# Assessing the potential cardiovascular risk of microdosing the psychedelic LSD in mice

**DOI:** 10.1101/2025.04.08.647757

**Authors:** Devin P. Effinger, Serena S. Schalk, Jillian L. King, Janae R. Strong, Caden K. O’Connell, Joselynn R. Calderon, John D. McCorvy, Scott M. Thompson

## Abstract

Microdosing, the prolonged ingestion of psychedelics at sub-hallucinogenic doses, has gained popularity for its perceived cognitive and emotional benefits. Psychedelics have high affinity for 5-HT_2B_ receptors, which cause heart disease with strong chronic activation. We investigated the effects of microdosed psychedelics on cardiovascular health in mice using electrocardiography after chronically administering either serotonin as a positive control or lysergic acid diethylamide (LSD) at two sub-hallucinogenic doses. Serotonin produced significant ventricular thickening at 4- and 8-weeks. No significant changes were observed in vehicle or LSD groups. We determined the affinity and potency of LSD, psilocybin, and norfenfluramine at mouse and human 5-HT_2B_Rs and observed no significant differences. We calculated that levels of 5-HT_2B_ activation by low-dose LSD were substantial, but short-lived, compared to the cardiotoxin *d*-fenfluramine. Together, these data provide no evidence of cardiovascular risk associated with prolonged administration of low-dose LSD in mice.

## Introduction

Psychedelic drugs, such as psilocybin and LSD, have produced therapeutic effects for a variety of affective psychiatric disorders.^1-3^ Colorado and Oregon recently passed legislation decriminalizing possession and use of psilocybin. There has subsequently been a 43% increase in past-year initiation of psychedelic use, versus a 15% increase in other US states. Many other states are considering legislation regarding medical use of psychedelic compounds.^4^ The public health consequences of this ‘experiment’ are likely to be many and multifaceted. Because so much remains unknown about long-term psychedelic use, it is a challenge to advise the public and potential users about many key issues surrounding safety.

Serotonergic psychedelics, such as psilocybin and LSD, are potent agonists at many serotonin receptors (5-HTRs), including the 5-HT_2A_ and 5-HT_2B_ receptors.^5,6^ Strong activation of 5-HT_2A_Rs causes the characteristic alterations in perception and consciousness.^7,8^ This psychedelic response is a potential barrier to widespread and cost-effective utilization of psychedelic compounds because it necessitates 6-8 hours of psychological support in an inpatient setting during administration^8^. For this reason, microdosing, or the chronic consumption of a sub-hallucinogenic dose of a psychedelic, typically LSD or psilocybin, has gained popularity due to anecdotal accounts of emotional and cognitive benefits. Between 2015 and 2023, internet searches related to microdosing increased by a factor of 13.4, with the greatest increases seen in Oregon and Colorado.^9^ A full understanding of the risks associated with chronic use of psychedelics is lacking, including cardiovascular risk.

Serotonergic appetite suppressants, such as *d*-fenfluramine, were associated with a significant incidence of valvulopathies and pulmonary hypertension, causing their withdrawal from the market.^10^ This pathology is characterized by echocardiographic evidence of heart wall thickening^11^ and mitral regurgitation, as well as histological evidence of mitral valve defects caused by cellular proliferation.^12^ 5-HT_2B_Rs are expressed at high levels in the embryonic and adult heart, where they are necessary for normal cardiac development,^13^ and their activation was subsequently identified as the cause of this cardiac valvulopathy and the pulmonary hypertension.^14^ Although the partially overlapping 5-HT_2B_R pharmacological profile of these medications with serotonergic psychedelics is known, the cardiovascular risks associated with use of psychedelics remain uncertain. While transient 5-HT_2B_R activation following a single acute administration of a high dose of psychedelics is unlikely to cause cardiac damage, it is not known whether longer-term consumption during microdosing causes damage. Given the increasing legalization, availability, and use of psychedelics, investigation into potential cardiovascular risks associated with prolonged psychedelic use is imperative.^15,16^

## Results

### Effects of LSD and 5-HT on head twitch response, heart rate, and body weight

In order to determine the mouse equivalent of a sub-hallucinogenic microdose, we used the head twitch response assay, as an indicator of 5-HT2AR activation.^17^ Male and female C57BL/6J mice were given intraperitoneal (i.p.) injections of LSD at 0.01 mg/kg, 0.03 mg/kg, or 0.1 mg/kg. Consistent with previous results,^18^ no significant increase in head twitch frequency was seen at 0.01 mg/kg and dose-dependent increases in head twitch frequency were seen at 0.03 and 0.1 mg/kg (**Fig. 1**). Use of 0.01 and 0.03 mg/kg are thus equivalent to a sub-hallucinogenic and weakly hallucinogenic dose, as might be used in human microdosing.

**Figure 1:**
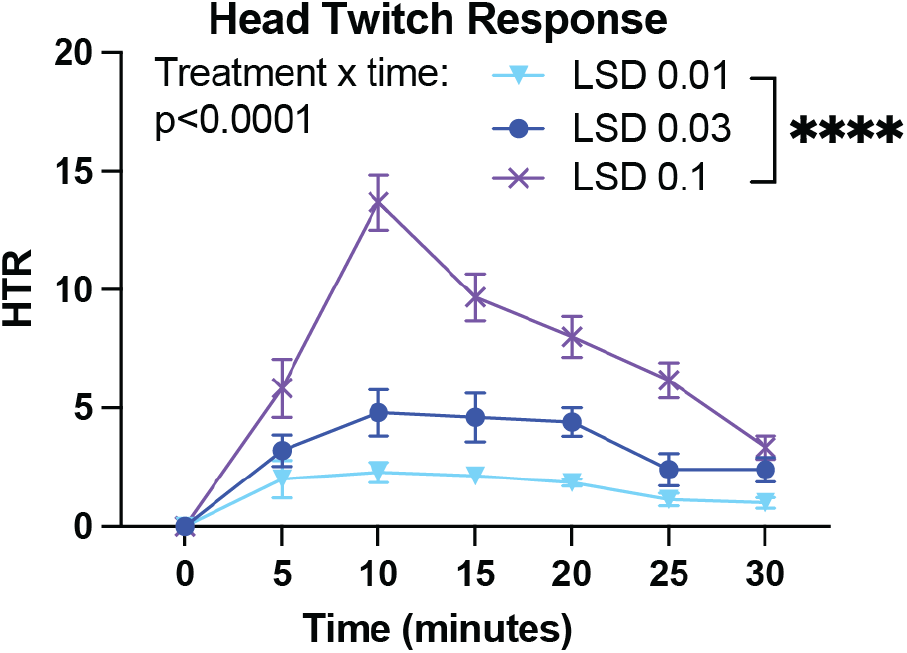
Head twitch response dose response assay to determine dosing. Head twitch response assay reveals that the two low doses of LSD used in the study (0.01 n=4m/3f and 0.03 mg/kg n=3m/2f) produce minimal increases in head twitch frequency, compared to the response elicited by a high dose of LSD (0.1 mg/kg n=3m/3f). 5-HT = serotonin, LSD = lysergic acid diethylamide.

We investigated the cardiovascular consequences of administering two different sub-hallucinogenic doses of LSD (0.01/0.03 mg/kg, i.p.) for eight weeks in mice, using a 5-days on/2 days off protocol, which is a widely used human microdosing paradigm^19^, as compared to a positive control group receiving injections of serotonin (40 mg/kg, i.p.), which has been shown to induce cardiac pathology in rodents.^20^ Echocardiography was performed at baseline, 4-weeks, and 8-weeks. There was no effect of treatment on heart rate or body weight during the treatment period, indicating no overall adverse effects of the treatment on animal health.

### Serotonin, but not LSD or vehicle, produced changes in echocardiographic structural and functional endpoints

To assess potential cell proliferative effects within the heart, left ventricle (LV) inner diameter, posterior wall diameter, and volume were measured. No significant change in LV inner diameter was seen in vehicle control mice for end diastolic or end systolic (**Fig. 2A**). In the 5-HT treated animals, there was a significant effect across time for end diastolic and systolic, with significant decreases in diameter compared to baseline seen at 4-weeks (**Fig. 2B**). No effects were seen in either the 0.01 or 0.03 mg/kg LSD groups (**Fig. 2C-D**).

**Figure 2:**
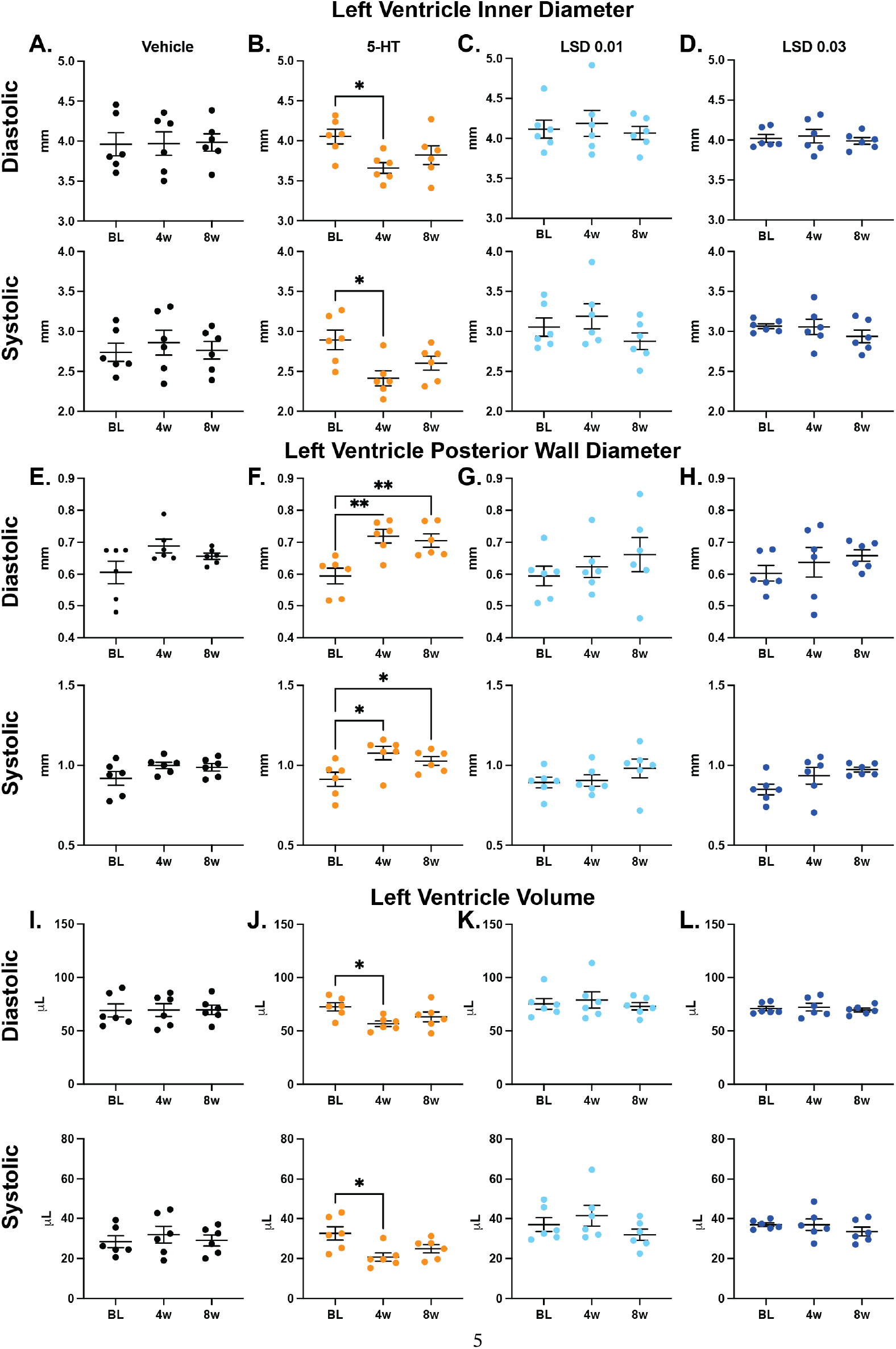
Serotonin, but not LSD or vehicle, produced changes in echocardiographic structural endpoints. *<u>LV inner diameter (n=6/group):</u>* **(A)** Vehicle end diastolic (top):; end systolic (bottom) **(B)** 5-HT end diastolic (top): 1-way ANOVA revealed an effect (F(1.761,8.807)=5.600, p=0.0296), Dunnett’s post hoc analysis found significant differences at 4-weeks (p=0235); end systolic (bottom): 1-way ANOVA revealed an effect (F(1.873,9.365)=4.915, p=0.0363), Dunnett’s post hoc analysis found significant differences at 4-weeks (p=0.0269). **(C)** LSD 0.01mg/kg end diastolic (top); end systolic (bottom) (**D)** LSD 0.03 mg/kg end diastolic (top); end systolic (bottom). *<u>LV posterior wall diameter (n=6/group):</u>* **(E)** Vehicle end diastolic (top); end systolic (bottom). **(F)** 5-HT end diastolic (top): 1-way ANOVA revealed an effect (F(1.984,9.919)=27.53, p<0.0001). Dunnett’s post hoc analysis found significant differences at 4- (p=0.0018) and 8-weeks (p=0.0038); end systolic (bottom): 1-way ANOVA revealed an effect (F(1.629,8.145)=8.971, p=0.0107). Dunnett’s post hoc analysis found significant differences at 4-(p=0.0319) and 8-weeks (p=0.0249). **(G)** LSD 0.01mg/kg end diastolic (top); end systolic (bottom). **(H)** LSD 0.03 mg/kg end diastolic (top); end systolic (bottom). *<u>LV volume (n=6/group):</u>* **(I)** Vehicle end diastolic (top); end systolic (bottom). **(J)** 5-HT end diastolic (top): 1-way ANOVA revealed an effect (F(1.705,8.526)=5.665, p=0.0305). Dunnett’s post hoc analysis found significant differences at 4-weeks (p=0.0241); end systolic (bottom): 1-way ANOVA revealed an effect (F(1.891,9.455)=4.853, p=0.0369). Dunnett’s post hoc analysis found significant differences at 4-weeks (p=0.0368). **(K)** LSD 0.01mg/kg end diastolic (top); end systolic (bottom). **(L)** LSD 0.03 mg/kg end diastolic (top); end systolic (bottom). BL = baseline, 4w = 4-week, 8w = 8-week, 5-HT = serotonin, LSD = lysergic acid diethylamide. \**p* < 0.05, \*\**p* < 0.01, *** *p* < 0.001, **** *p* < 0.0001.

For LV posterior wall diameter, no effect was seen in the vehicle control group (**Fig. 2E**). In the 5-HT group, there was a significant effect across time, with significant increases in wall diameter at 4- and 8-weeks compared to baseline (**Fig. 2F**). No effect was seen in either the 0.01 or 0.03 mg/kg LSD groups (**Fig. 2G-H**).

For LV volume, no effect was seen in the vehicle control group (**Fig. 2I**). In the 5-HT group, there was a significant effect across time for both end diastolic and systolic, with significant decreases in volume seen at 4-weeks compared to baseline (**Fig. 2J**). No effect was seen in either the 0.01 or 0.03 mg/kg LSD groups (**Fig. 2K-L**).

To assess functional consequences, analysis of ejection fraction was conducted. No changes were detected in the vehicle or 5-HT control groups (**Fig. 3A-B**). In the 0.01 mg/kg LSD group, there was an effect across time, however, post hoc analysis revealed no differences from baseline at follow up scans (**Fig. 3C**). In the LSD 0.03 mg/kg group, no changes were detected (**Fig. 3D**). Next, fractional shortening was assessed. No changes were detected in the vehicle and 5-HT control groups (**Fig. 3E-F**). In the 0.01 mg/kg LSD group, there was an effect of time, however, post hoc analysis found no differences between baseline or follow up time points (**Fig. 3G**). In the 0.03 LSD mg/kg group, there were no changes in fractional shortening or ejection fraction (**Fig. 3H**).

**Figure 3:**
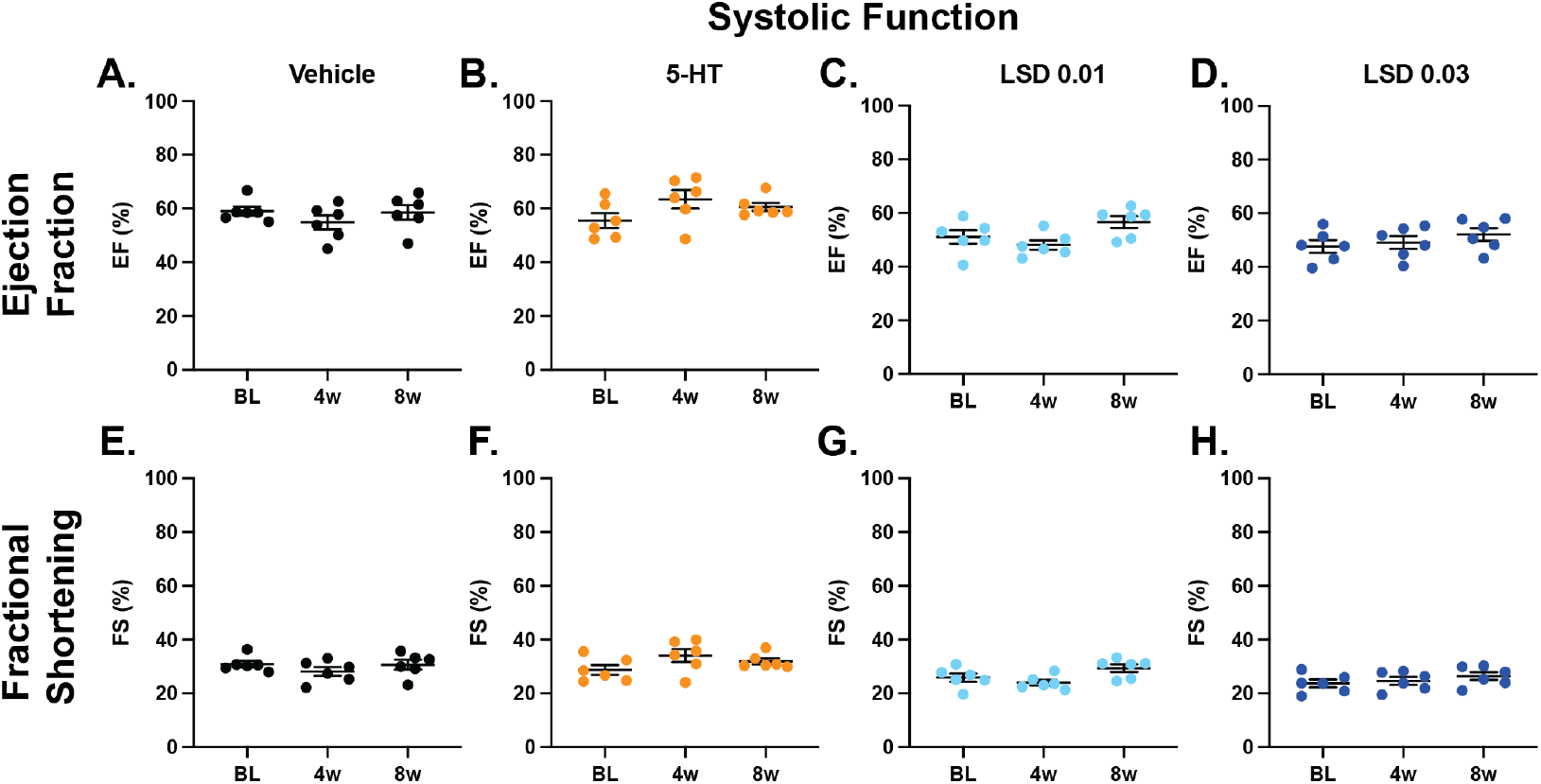
Serotonin, LSD, and vehicle produced no changes in echocardiographic functional endpoints. <u>Ejection fraction</u>*<u>(n=6/group)</u>*: (A) vehicle. (B) 5-HT. (C) LSD 0.01 mg/kg: 1-way ANOVA found a significant effect across time (F(1.705,8.526)=5.665, p=0.0305). Dunnett’s post hoc analysis revealed no significant differences at follow up time points compared to baseline. (D) LSD 0.03 mg/kg. <u>Fractional shortening</u>*<u>(n=6/group</u>)*: (E) vehicle. (F) 5-HT. (G) LSD 0.01 mg/kg: 1-way ANOVA found a significant effect (F(1.766,8.829)=5.367, p=0.0326). Dunnett’s post hoc analysis revealed no significant differences at follow up time points compared to baseline. (H) LSD 0.03 mg/kg. BL = baseline, 4w = 4-week, 8w = 8-week, 5-HT = serotonin, LSD = lysergic acid diethylamide, EF = ejection fraction, FS = fractional shortening. \**p* < 0.05, \*\**p* < 0.01, *** *p* < 0.001, **** *p* < 0.0001.

### Comparison of agonist binding at mouse and human 5-HT_2B_ receptors and estimation of 5-HT_2B_ receptor occupancy after microdoses

LSD is an ergoline psychedelic and affinity values between human and mouse 5-HT_2B_Rs may differ by more than an order of magnitude for ergolines such as methysergide and mesulergine.^21^ We therefore used a previously described bioluminescence resonance energy transfer (BRET) assay^6^ to measure G_q_ dissociation activity for 5-HT, LSD, *d*-norfenfluramine, and psilocin at both human and mouse 5-HT_2B_Rs. Only small differences in potency (EC_50_) and efficacy (E_max_) were observed between species (**Fig. 4A-D, Supplement Table 1**).

**Figure 4:**
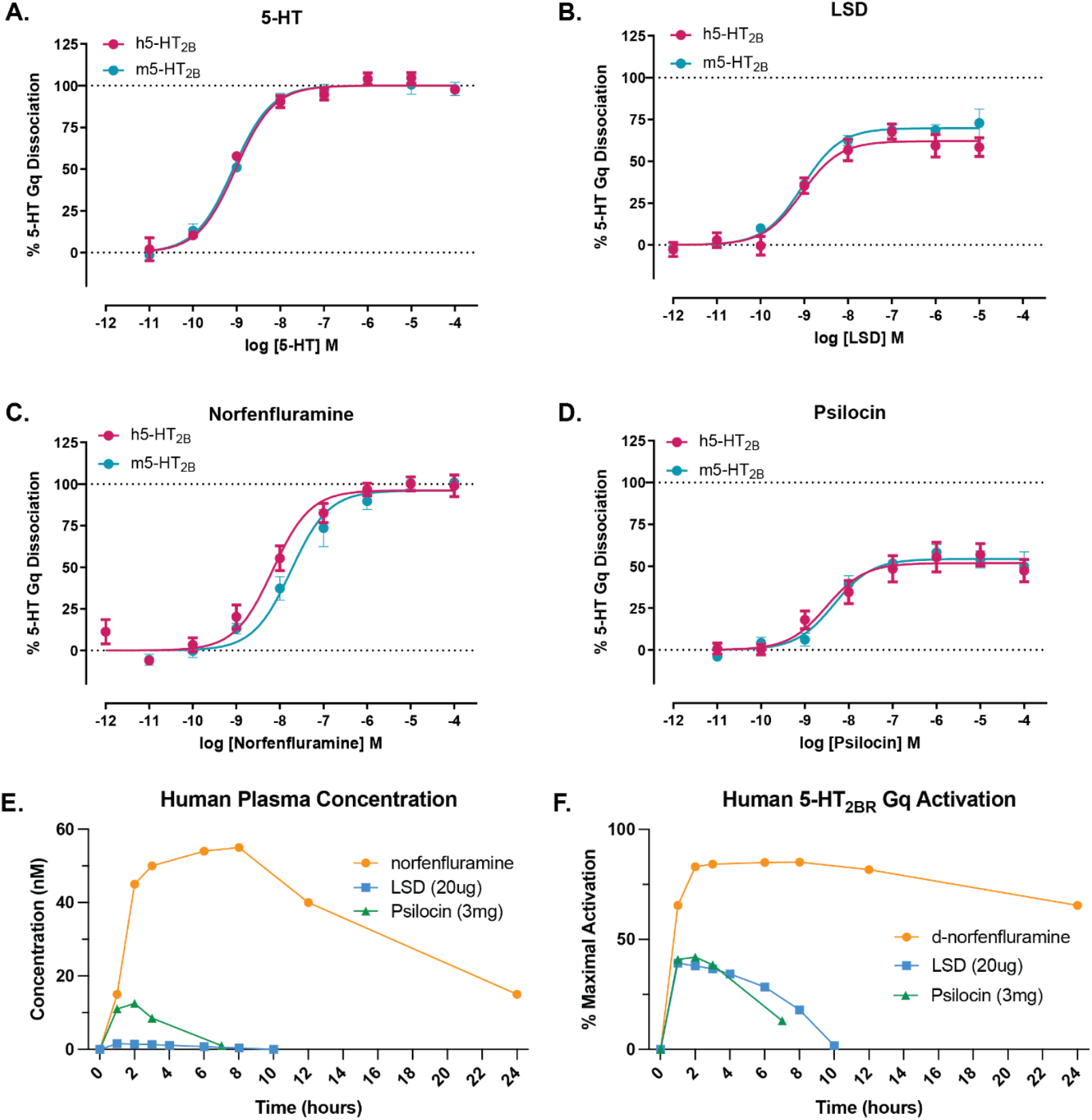
Comparison of agonist binding at mouse and human 5-HT_2B_ receptors and estimation of 5-HT_2B_ receptor occupancy after microdoses. Dose-response relationships for activation of Gq dissociation at recombinant mouse (blue) and human (red) 5-HT_2B_Rs for 5-HT **(A)**, LSD **(B)**, *d-*norfenfluramine **(C)**, and psilocin **(D)**. Data represent mean ± SEM for three independent experiments. **(E)** The time course of plasma levels of microdoses of LSD (20ug, blue) or psilocin (3mg, green), and a full dose of *d*-norfenfluramine (30mg, orange), as taken from published literature. **(F)** Calculated time course of the percent of maximal 5-HT_2B_R-activation produced by the plasma profiles of LSD, psilocin, and norfenfluramine shown in **E**.

To better understand why low-dose LSD did not induce 5-HT_2B_R-mediated valvulopathy, we estimated the fraction of 5-HT_2B_R activation activated by a single human microdose of LSD (20 µg) and compared it to the 5-HT_2B_R activation expected from a single human dose *d*-fenfluramine (30 mg) or a microdose of psilocybin (3 mg). The time course of the plasma levels of LSD, the active *d*-fenfluramine metabolite, *d*-norfenfluramine, and the active psilocybin metabolite, psilocin, were taken from published papers.^22,23,24^ Fractional receptor activation was then computed from the Hill equation, using EC_50_ aqnd E_max_ values for 5-HT_2B_ activation determined above. Consistent with the larger dose, the pharmacokinetic profile of *d*-norfenfluramine was considerably larger and longer than for LSD (**Fig. 4E**).

Comparing 5-HT_2B_R activation across time, we found that 5-HT_2B_R occupancy for microdoses of LSD and psilocybin peaked at about half the level achieved with norfenfluramine consumption and was absent by 10 hours, whereas *d*-norfenfluramine-induced 5-HT_2B_R activation remained >50% for 24 hours (**Fig. 4F**). Given that fenfluramine was a daily prescribed treatment, it is likely that 5-HT_2B_R activation would be constant and increasing, which could therefore explain its greater cardiotoxic effects. Together, these data suggest that the risk of LSD- or psilocybin-induced cardiotoxicity is lower than for *d*-fenfluramine due to their lower concentration and shorter half-life in plasma.

**Supplementary Table 1.**
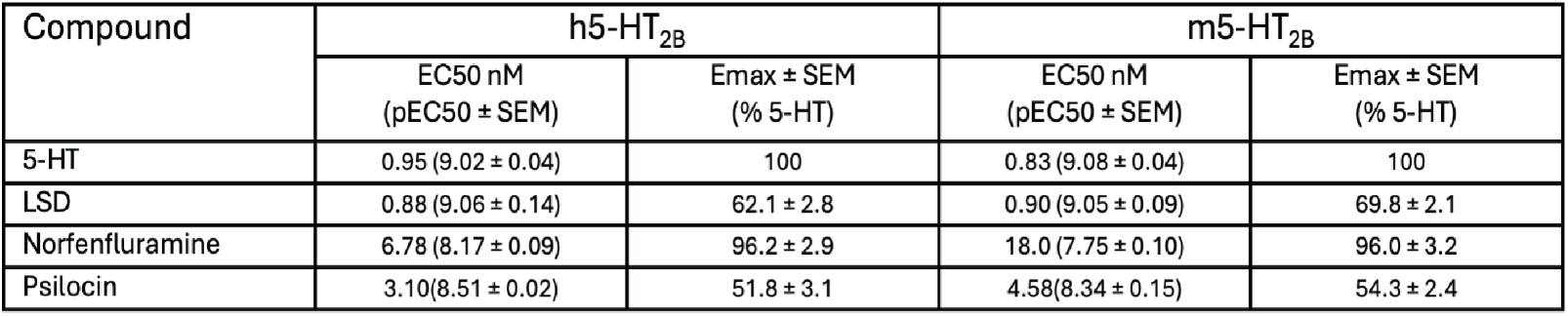
EC_50_ and E_max_ values from 5-HT_2B_R Gq dissociation assays.

| Compound | h5-HT<sub>2B</sub> |  | m5-HT<sub>2B</sub> |  |
| --- | --- | --- | --- | --- |
|  | EC50 nM<br>(pEC50 ± SEM) | E<sub>max</sub> ± SEM<br>(% 5-HT) | EC50 nM<br>(pEC50 ± SEM) | E<sub>max</sub> ± SEM<br>(% 5-HT) |
| 5-HT | 0.95 (9.02 ± 0.04) | 100 | 0.83 (9.08 ± 0.04) | 100 |
| LSD | 0.88 (9.06 ± 0.14) | 62.1 ± 2.8 | 0.90 (9.05 ± 0.09) | 69.8 ± 2.1 |
| Norfenfluramine | 6.78 (8.17 ± 0.09) | 96.2 ± 2.9 | 18.0 (7.75 ± 0.10) | 96.0 ± 3.2 |
| Psilocin | 3.10(8.51 ± 0.02) | 51.8 ± 3.1 | 4.58(8.34 ± 0.15) | 54.3 ± 2.4 |

## Discussion

Given that serotonergic psychedelics, including psilocybin and LSD, are potent agonists of the 5-HT_2B_R, like the known cardiotoxic metabolite *d*-fenfluramine, there is concern about the potential for use of these compounds to have deleterious cardiotoxic effects. Although this seems unlikely following a single high-dose administration, the concerns are greater for chronic consumption of psychedelics, even with small microdoses.^15,16^ We used echocardiography in mice to assess the potential cardiovascular risk associated with prolonged administration of two low-doses (0.01 and 0.03 mg/kg, i.p.) of the psychedelic LSD, compared to prolonged dosing of 5-HT (40mg/kg, i.p.) as a positive control.^20^ As predicted, the 5-HT-treated group had a significant thickening of the left ventricular wall and decreased ventricular volume. However, no changes in fractional shortening or ejection fraction were found. These results are consistent with studies showing that overexpression of 5-HT_2B_Rs in mice leads to increases in ventricular wall thickness, accompanied by increased cellular proliferation, without changes in ventricular volume, fractional shortening, blood pressure, or heart rate; presumably due to compensatory remodeling.^25^ These results are also comparable to pathological changes observed in humans following chronic use of *d-* fenfluramine and other serotonergic appetite suppressants.^10^

No significant effects of treatment on ventricular inner diameter, posterior wall thickness, or ventricular volume were seen after injections of saline or in either low-dose LSD group at 4- or 8-weeks post-treatment. Finally, chronic administration of neither 5-HT nor LSD produced differences in body weight or heart rate across the 8-week time frame.

Given the high affinity of LSD for 5-HT_2B_Rs, why did 8-weeks of chronic administration fail to induce cardiovascular pathology? One potential explanation is that the treatment period was not long enough. Given the pathological changes observed with 4-to 8-weeks of 5-HT administration, chronic activation of 5-HT_2B_Rs over this time-period would seem sufficiently long to detect pathological changes if they were triggered by microdosed LSD. An alternative explanation is that the pharmacokinetics of LSD at these doses are insufficient to produce significant 5-HT_2B_R activation. The two doses of LSD used here are presumably comparable to the non-hallucinogenic doses used in human microdosing. The lowest dose (0.01 mg/kg) failed to produce a head twitch response in mice, a behavioral readout associated with strong 5-HT_2A_ receptor activation by classical psychedelic drugs,^17^ and the 0.03 mg/kg dose produced only a small response.

5-HT_2B_R-activation of G_q_ signaling-initiated second messenger pathways was idenitifed as the cause of the pro-proliferative, cardiotoxic effects of fenfluramine.^26,27^ We determined the potency and efficacy of LSD, psilocybin, and norfenfluramine to activate 5-HT_2B_R-dependent signaling via a G_q_ dissociation activity assay at both human and mouse receptors. Observed differences in affinity between norfenfluramine (EC_50_ = 6.78nM), LSD (0.88nM), and psilocin (3.1nM) were small, but there were larger differences in potency, with norfenfluramine capable of 96% of the activation produced by 5-HT, but LSD and psilocin only capable of 62% and 52% of the activation produced by 5-HT. These data were then combined with previously published plasma concentration data in humans to estimate the fraction of available 5-HT_2B_Rs activated. A single dose of *d*-fenfluramine results in plasma concentrations of its active metabolite that are significantly higher and more prolonged than plasma levels of LSD or psilocin following a low dose of LSD or psilocybin. The fraction of 5-HT_2B_R activation was estimated to be sizable for all three compounds, but was greater and more prolonged for *d-*fenfluramine than for LSD or psilocybin. We thus attribute the absence of cardiovascular pathology after microdoses of LSD in our experiments to insufficient levels of 5-HT_2B_R receptor occupancy and concomitant of activation of G_q_ signaling. It should be noted that MDMA may also elevate risk for cardiac disease through these same mechanisms.^28^

An essential caveat of this preclinical work is that although we have no evidence that microdosing LSD results in cardiac pathology in mice, we cannot therefore conclude that the human microdosing of LSD is safe. While there is evidence that the 5-HT_2B_R has species specific binding properties,^20^ we did not detect any species differences in affinity or efficacy for LSD or psilocybin in 5-HT_2B_ receptor stimulated G_q_ dissociation. This finding does not preclude that there may also be significant differences in the rate at which psychedelics are absorbed or metabolized in mice versus humans, thus changing the degree of 5-HT_2B_ receptor occupancy. Furthermore, our studies were performed on presumably healthy mice and the risks for significant cardiotoxicity from microdosed psychedelics could be greater in people with pre-existing cardiovascular risk factors. Prospective studies should be performed to determine whether microdosed psychedelics produce evidence of cardiac pathology in humans.

## Methods

### Animals

All procedures were approved by the Institutional Animal Care and Use Committee (IACUC) at the University of Colorado Anschutz Medical Campus. Male and female C57BL/6J mice were obtained from Jackson Laboratories and ordered to arrive at 8-weeks old. Animals were allowed to habituate for 1-week prior to initiating echocardiograms.

### Drugs

For cardiovascular endpoints, serotonin hydrochloride (Tocris Bioscience) was dissolved in saline and injected at 40 mg/kg (i.p.). The serotonin dose was chosen based on the study of Gustafsson et al.,^19^ which demonstrated evidence of cardiac pathology. (+)-Lysergic acid diethylamide tartrate was provided by the National Institute on Drug Abuse Drug Supply Program (NDSP) and was dissolved into saline prior to being injected at 0.01, 0.03, or 0.1 mg/kg (i.p.). For pharmacological endpoints, 5-HT and (+)-norfenfluramine were purchased from Sigma-Aldrich, and LSD and psilocin were purchased from Cayman Chemical Company.

### Head Twitch Response

Mice were recorded with a high-speed video camera and head twitches was scored manually by a blinded observer.

### Rodent echocardiograms

Echocardiographic recordings were conducted using FujiFilm Visualsonics F2 Imaging system and analyzed using VevoLab Imaging Analysis software. Animals were first weighed and then anesthetized with an initial bolus of 3% isoflurane and then maintained with 1.5% isoflurane during the echo procedure. The animals were laid supine on a warming pad to maintain body temperature at 37 degrees Celsius; temperature was monitored and controlled with a rectal thermometer in series with a heated platform throughout the experiments. A thin layer of Nair was placed on the skin of the chest for 15 seconds. Wet gauze was used to wipe away the Nair and hair from the chest. The chest was then covered with warm ultrasound gel. Echocardiographic images were obtained while the echo probe was held gently against the chest. The echocardiography procedure lasted between 15 – 30 minutes, depending on the number of individual images obtained. At the end of the procedure, the ultrasound gel was removed with wipes and water. Animals were allowed to recover on a table by the echo instrument in a cage placed over a warming pad before returning to the housing room.

### 5-HT_2B_R Gq dissociation BRET assays

Bioluminescence resonance energy transfer G protein dissociation assays (BRET) measuring 5-HT_2B_ Gq and β3/γ9 dissociation were conducted in HEKT cells (ATCC CRL-11269; mycoplasma-free). Cells were transfected and compounds were incubated with cells for 60 minutes at 37°C and read on a PheraStarFSX (BMG LabTech), as described previously.^6^ Human and mouse 5-HT_2B_R constructs were subcloned into vectors, as described previously.^6^ Data were analyzed using nonlinear regression “log(agonist) vs. response” in Graphpad Prism 10 to yield Emax and EC50 parameter estimates. Data were normalized to percent 5-HT stimulation, with a concentration–response curve was present on every plate.

### Pharmacokinetic Modeling

The time course of human plasma drug levels was used to generate 5-HT2BR activation profiles for LSD, psilocybin, and norfenfluramine using the Hill equation:

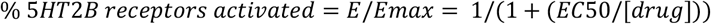

where EC50 is the concentration producing 50% receptor activation.

### Statistical Analysis

1-way analysis of variance (ANOVA) tests were performed in GraphPad Prism Version 10.4.1. Upon finding of significant main effect and/or interactions, Dunnett’s post hoc multiple comparisons were done to compare follow up time points to baseline recording. Statistical details, including F values and n can be found in figure legends. 2-way ANOVA were performed to assess any changes in body weight or heart across time and between conditions.

## Resource availability

### Lead Contact

- Requests for further information and resources should be directed to and will be fulfilled by the lead contact, Scott Thompson.

### Materials availability

- This study did not generate new unique reagents.

### Data and Code Availability

- All data reported in this paper will be shared by the lead contact upon request.

## Acknowledgements

This research was supported by National Institutes of Health Grant R01 DA 061433 (to JDM) and the CU Anschutz Department of Psychiatry (SMT). We would like to thank the National Institute on Drug Abuse Drug supply program for graciously providing the LSD tartrate. Additionally, we would like to acknowledge the Preclinical Cardiovascular Core at CU Anschutz for their performance and analysis of echocardiograms.

## Author Contributions

D.P.E conception, data collection, drug administration, data analysis, manuscript drafting, manuscript editing; S.S.S. data collection; J.L.K. drug administration, manuscript editing; J.R.S. drug administration, data collection; C.K.O. data analysis; J.R.C. drug administration; J.D.M. funding, manuscript editing; S.M.T. funding, data analysis, manuscript drafting, manuscript editing.

